# Telomere replication stress-induced DNA damage response triggers inflammatory signaling via canonical and non-canonical STING pathways

**DOI:** 10.1101/2025.07.11.664434

**Authors:** Wei Zhu, Yi Gong, Yunong Wang, Myriam Gorospe, Yie Liu

**Affiliations:** Laboratory of Genetics and Genomics, National Institute on Aging/National Institutes of Health, 251 Bayview Blvd, Baltimore, MD 21224, USA

**Keywords:** Telomere replication, telomere fragility, DNA damage response, TRF1, cytosolic DNA, cGAS, STING, inflammatory response

## Abstract

Telomeres are protected by the shelterin complex, but they are also common fragile sites and are particularly susceptible to replicative stress. We found that depletion of telomeric repeat-binding factor 1 (TRF1), a key shelterin component essential for telomere replication, in mouse embryonic fibroblasts (MEFs) activated ATR- and subsequent ATM-dependent DNA damage responses. TRF1 loss increased the formation of micronuclei and cytosolic DNA, leading to ATR-dependent micronuclear rupture and activation of the cGAS/STING pathway. ATM activation enhanced STING K63 modification, thereby boosting the STING/NFκB pathway. Inhibition of ATM or cGAS reduced the expression of the pro-inflammatory cytokine IL6, with combined inhibition further suppressing IL6 levels. Depletion or inhibition of STING alone decreased production of IL6 and IFNβ, with no major reduction by combined ATM and/or cGAS inhibitors. These findings indicate that STING acts epistatically with ATM- and cGAS-mediated inflammatory responses. Overall, the telomere replication stress and dysfunction triggered by loss of TRF1 promotes inflammation through the ATR/cGAS/STING and ATM/STING pathways.

## INTRODUCTION

Telomeres are nucleoprotein structures at the ends of chromosomes, consisting of repetitive TTAGGG DNA sequences and shelterin protein complexes. They function to cap the chromosome ends from triggering a DNA damage response (DDR). Disruption of the protective cap can trigger DDR, leading to cellular senescence and the development of a senescence-associated secretory phenotype (SASP) [1–6]. Telomeres are also susceptible to the formation of alternative structures, such as G-quadplexes, which can induce replication fork stress and breakage [7, 8]. Thus, telomeres exhibit similar characteristics as common genomic fragile sites, which are hypersensitive to replication stress, pose challenges to the DNA replication machinery, and are hotspots for chromosome breakage [9–11].

Shelterin is crucial for maintaining telomere structure, capping function, and overall stability [12]. The key shelterin protein, TRF1, directly binds to double-stranded telomeric DNA and plays a vital role in telomere semi-conservative DNA replication and capping. TRF1 facilitates the resolution of G-quadruplex DNA structures, thereby promoting efficient telomere replication and preventing replication fork stalling [9, 11–13]. In contrast, loss of TRF1 impairs telomere replication fork progression, increases telomere fragility and the formation of telomere damage foci, activates the DDR, and ultimately induces cellular senescence [10, 11].

The cGAS-STING pathway is essential for linking DNA damage to the inflammatory response [14]. Defects in DNA replication can generate extra-genomic DNA fragments, particularly when the replication process is disrupted [15]. The release of genomic DNA from stalled replication forks or incomplete replication results in the accumulation of micronuclei, single-stranded DNA (ssDNA), or double-stranded DNA (dsDNA) in the cytosol [15–17]. DNA fragments sequestered within micronuclei can be further released into the cytosol upon rupture of the fragile micronuclear envelope [15]. As a key component of the innate immune response, the cGAS-STING pathway is particularly important for sensing cytosolic DNA [18]. Cyclic GMP-AMP synthase (cGAS) functions as a cytosolic DNA sensor [18, 19]. It binds to naked cytosolic DNA and synthesizes cyclic GMP-AMP (cGAMP), a second messenger that induces oligomerization and activation of stimulator of interferon genes (STING) in the endoplasmic reticulum (ER) and facilitates its exit [20–22]. Activated STING recruits TANK-binding kinase 1 (TBK1), triggering its autophosphorylation and subsequent recruitment and activation of interferon regulatory factor 3 (IRF3) and inhibitor of κB kinase (IKK) [21, 23–26]. Phosphorylated IRF3 dimerizes and translocates to the nucleus, where it induces the transcription of type I interferons (IFNs) and other interferon-stimulated genes [22, 24, 25, 27]. On the other hand, the STING-TBK1 complex activates IKK, leading to phosphorylation and degradation of IκB, which releases nuclear factor kappa B (NFκB) for nuclear translocation and promotes transcription of target genes [21, 28, 29]. In addition to the canonical cGAS-STING pathway, STING can also be activated through noncanonical mechanisms independent of cGAS, such as DNA damage [30–32], infection with enveloped RNA viruses [33], and specific protein-protein interactions [34]. Notably, previous studies have shown that ATM promotes K63-linked ubiquitination of STING and preferentially activates the NFκB pathway [30–32]. Thus, both canonical and noncanonical STING pathways serve as critical links between DNA damage and the inflammatory response [16, 18, 32, 35].

In this study, we utilized TRF1-deficient mouse embryonic fibroblasts (MEFs) to investigate the mechanistic basis of telomere replication stress and telomere dysfunction-induced DDR in inflammation response. We found that loss of TRF1 activated ATR, followed by subsequent activation of ATM. TRF1-deficient MEFs exhibited increased formation of micronuclei and cytosolic telomeric DNA fragments, alongside activation of the cGAS/STING pathway.

Furthermore, TRF1 loss resulted in synergistic regulation of inflammation through both the ATM/STING and ATR/cGAS/STING axes, which was mitigated by STING depletion or inhibition. Collectively, our findings highlight the central role of STING in mediating the interplay between telomere replication stress and inflammatory responses.

## RESULTS

### Loss of TRF1 induces the formation of cytosolic and micronuclei DNA fragments

TRF1 is essential for telomere capping and efficient telomere replication [12]. Previous studies have shown that deletion of TRF1 triggers the DDR. Sfeir et al. demonstrated that TRF1 loss activates the ATR pathway, while Martínez et al. reported stimulation of the ATM pathway. Both findings demonstrated that TRF1 deficiency elevates telomere replication stress, leading to increased telomere fragility and the formation of DNA damage foci [10, 11]. To investigate whether telomere fragility and damage contribute to the accumulation of DNA fragments in the cytoplasm and micronuclei, we utilized TRF1^F/F^ Rosa26Cre-ER^T1^ MEFs [11]. Western blot (Fig S1A) and RT-qPCR (Figure S1B) analyses confirmed an efficient TRF1 deletion following Cre-mediated excision induced by OHT treatment. Consistent with previous reports, TRF1 deletion activated the DDR, as evidenced by increased p21 expression and the formation of 53BP1 foci at both genome-wide and telomeric sites (Figure S1B-E). In addition, the ATR pathway was activated, as evidenced by the elevated levels of phosphorylated ATR (p-ATR) and CHK1 (p-CHK1) 24 hours after OHT treatment (Figure 1A. Notably, ATM pathway activation occurred slightly later, with significant increases in phosphorylated ATM (p-ATM) and CHK2 (p-CHK2) detected 48 hours post-treatment (Figure 1B). Telomere fluorescence in situ hybridization (FISH) analysis further revealed fragile telomeres, signal-free ends, and end-to-end fusions in metaphase spreads of TRF1-deficient cells (Figure S1F), indicating compromised telomere replication and genomic integrity.

**Figure 1:**
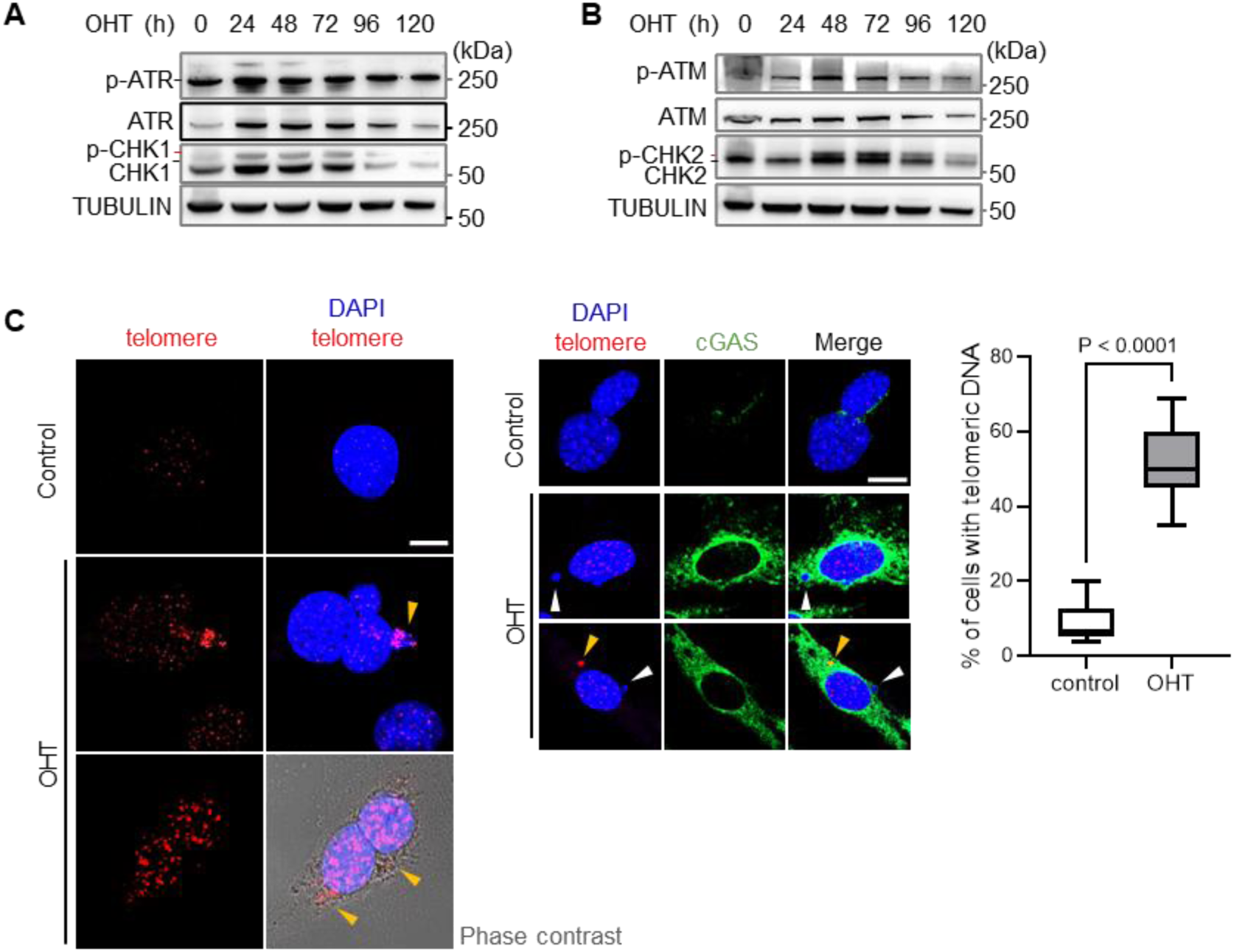
TRF1 depletion activates ATR- and ATM-mediated DDR and leads to the formation of extratelomeric DNA fragments in the cytoplasm and micronuclei in MEFs. (A, B) TRF1 depletion induces ATR (A) and ATM (B) activation in MEFs treated with OHT for 0–120 hours, as shown by Western blot analysis with the indicated antibodies. Black dashes on the left indicate the position of the protein of interest; red dashes indicate phosphorylated proteins. (C) Detection of DNA fragments in the cytoplasm or micronuclei that are either positive (yellow arrows) or negative (white arrows) for telomere signals, as determined by telomere FISH using a telC-Cy3 probe in MEFs treated with OHT (left panel). Phase contrast images highlight cytoplasmic telomeric signals. Immunofluorescence analysis of cGAS (green) is shown for MEFs under mock control (ctr) and OHT (+) treatment (middle panel). Scale bar, 10 μm. Quantification (right panel) shows the percentage of cells with telomeric signals in the cytoplasm or micronuclei. Approximately 100 cells were analyzed per group. P values were calculated using Student’s t-test. Data are presented as mean ± SD.

To determine whether damaged telomeric DNA is transferred to the cytoplasm and micronuclei, we performed telomere FISH to detect telomeric DNA in these cellular localizations. Following TRF1 deletion, we observed a significant increase in telomeric signals within both the cytoplasm and micronuclei (Figures 1C). Additionally, micronuclei lacking telomeric signals were also present in TRF1-deficient cells (Figure 1C). Similar features were observed in wild-type MEFs subjected to replication stress with 0.2 µM Aphidicolin treatment (Figure S1G-H). These results indicate that deficiencies in telomere replication and capping lead to DNA fragility and damage, resulting in the release of DNA fragments, including telomeric DNA, into the cytosol and micronuclei.

### The cGAS/STING pathway is activated as a result of TRF1 depletion and ATR activation

We then investigated whether the cytoplasmic DNA fragments and micronuclei in TRF1-deficient MEFs could activate the cGAS/STING pathway. Immunofluorescence analysis revealed increased cytosolic cGAS expression in OHT-treated MEFs (Figure 1C, middle panel). Western blot analysis demonstrated a transient elevation in phosphorylated STING (p-STING) levels following OHT treatment (Figure 2A). In addition, the levels of 2’3’-cGAMP, the product of cGAS activation by cytosolic DNA fragments, were elevated in OHT-treated cells (Figure 2B). We also observed STING aggregation in the ER, consistent with its oligomerization upon activation (Figures 2C). Collectively, these results demonstrate that cytosolic DNA generated by TRF1 depletion activates the cGAS/STING pathway.

**Figure 2:**
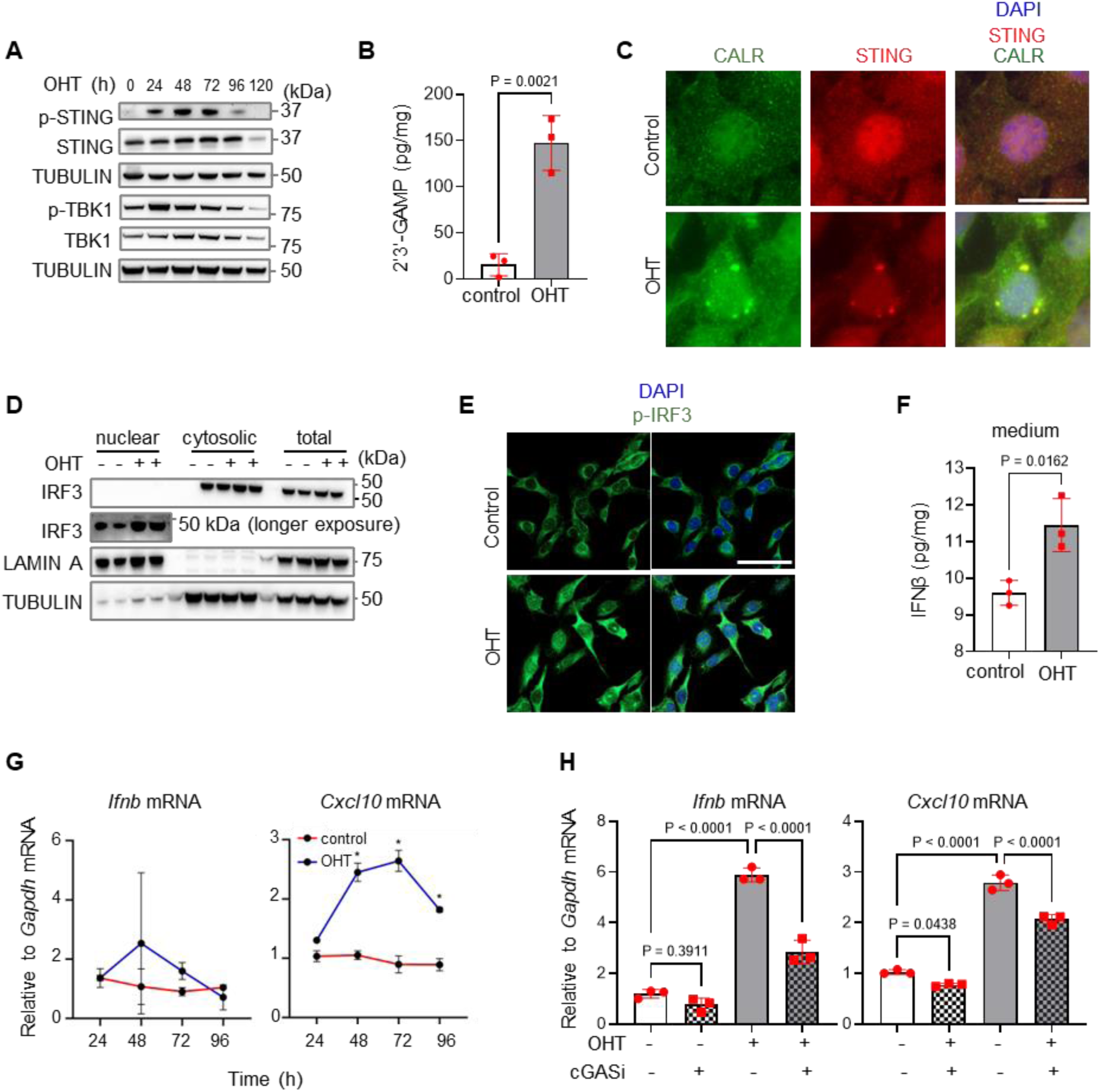
TRF1 depletion activates the cGAS/STING/IRF3 pathway. (A) Western blot analysis of cGAS/STING pathway proteins in MEFs treated with OHT (+) for the indicated durations. (B) Quantification of 2’3’-cGAMP levels in MEFs following mock (ctr) or OHT (+) treatment, measured by ELISA and normalized to total protein concentration in the cell pellet. (C) Immunofluorescence analysis of STING (red) and the ER marker calreticulin (CALR, green) in MEFs with mock (ctr) or OHT (+) treatment. Colocalization of STING and calreticulin signals is shown in an OHT-treated MEF. Nuclei are stained with DAPI (blue). Scale bar, 10 μm. (D) Western blot analysis of IRF3 in nuclear and cytosolic fractions and whole cell lysates from MEFs treated with mock (-) or OHT (+). A longer exposure of IRF3 is shown (lower panel). (E) Immunofluorescence analysis of p-IRF3 (green) indicating nuclear translocation in OHT-treated MEFs. DAPI is shown in blue. Scale bar, 50 μm. (F) Measurement of IFNβ protein levels in culture medium, normalized to total protein concentration in the cell pellet. (G-H) Relative mRNA levels of *Ifnb* and its inducible gene *Cxcl10* were determined by RT-qPCR: (G) in MEFs treated with mock (ctr) or OHT for the indicated times; (H) in MEFs treated with OHT and subsequently with cGAS inhibitor (cGASi) for 48 hours. P values were calculated using Student’s t-test (B, F) or one-way (H) and two-way (G) ANOVA for multiple comparisons. Data are presented as mean ± SD from three independent experiments for panels B and F, and as representative data from one of three independent biological experiments for panels G and H.

Activation of the cGAS/STING pathway leads to the phosphorylation and activation of TBK1, which subsequently phosphorylates IRF3, enabling its translocation to the nucleus. Once in the nucleus, IRF3 primarily promotes the transcription of interferon-stimulated genes, including IFNβ and related response elements such as CXCL10 [32, 36–38]. We therefore investigated whether TRF1 depletion elicits similar activation of the TBK1/IRF3/IFNβ axis. Western blot analysis revealed increased levels of phosphorylated TBK1 (p-TBK1) following OHT treatment (Figure 2A). Elevated IRF3 was detected in the nuclear fraction of OHT-treated MEFs (Figure 2D), and immunofluorescence analysis demonstrated increased nuclear staining of phosphorylated IRF3 (p-IRF3) after OHT treatment (Figure 2E). These results indicate that TRF1 deletion promotes the nuclear translocation of p-IRF3, a hallmark of IRF3 activation.

To assess IFNβ activation we measured the dynamic expression of IFNβ and its target gene CXCL10 following OHT treatment. ELISA showed upregulation of IFNβ protein levels in the culture medium post-OHT treatment (Figure 2F). Concurrently, RT-qPCR analysis revealed a moderate increase in *Ifnb* mRNA expression over a short time frame, while *Cxcl10* mRNA was readily detected at various time points (Figure 2G). To further confirm the role of cGAS in the induction of interferon-stimulated genes, we treated both mock and OHT-treated MEFs with the cGAS inhibitor RU.521. Subsequent RT-qPCR analysis demonstrated that RU.521 attenuated the OHT-induced levels of *Ifnb* and *Cxcl10* mRNAs (Figure 2H).

ATR activation has been reported to facilitate micronuclei rupture, thereby triggering the cGAS/STING pathway [39]. In our study, we observed activation of the ATR pathway 24 hours after OHT treatment (Figure 1A). To elucidate the role of ATR in cGAS/STING pathway activation in TRF1-deficient cells, we treated OHT-exposed MEFs with the ATR inhibitor (ATRi) VE-821.

The effectiveness of VE-821 in suppressing ATR activity was confirmed by reduced phosphorylation of CHK1 (Figure S2A). Notably, VE-821 treatment increased the number of micronuclei per cell in TRF1-deficient MEFs compared to mock-treated controls (Figure 3A). However, levels of 2’3’-cGAMP and IFNβ protein in the culture medium were reduced upon VE-821 treatment (Figures 3B and 3C). Furthermore, VE-821 treatment resulted in a more pronounced decrease in the levels of *Ifnb* mRNA compared to *Il6* mRNA (Figure 3D). These findings suggest that ATR activation promotes micronuclei rupture following TRF1 deletion, which in turn activates the cGAS/STING pathway and preferentially enhances the IFNβ response.

**Figure 3:**
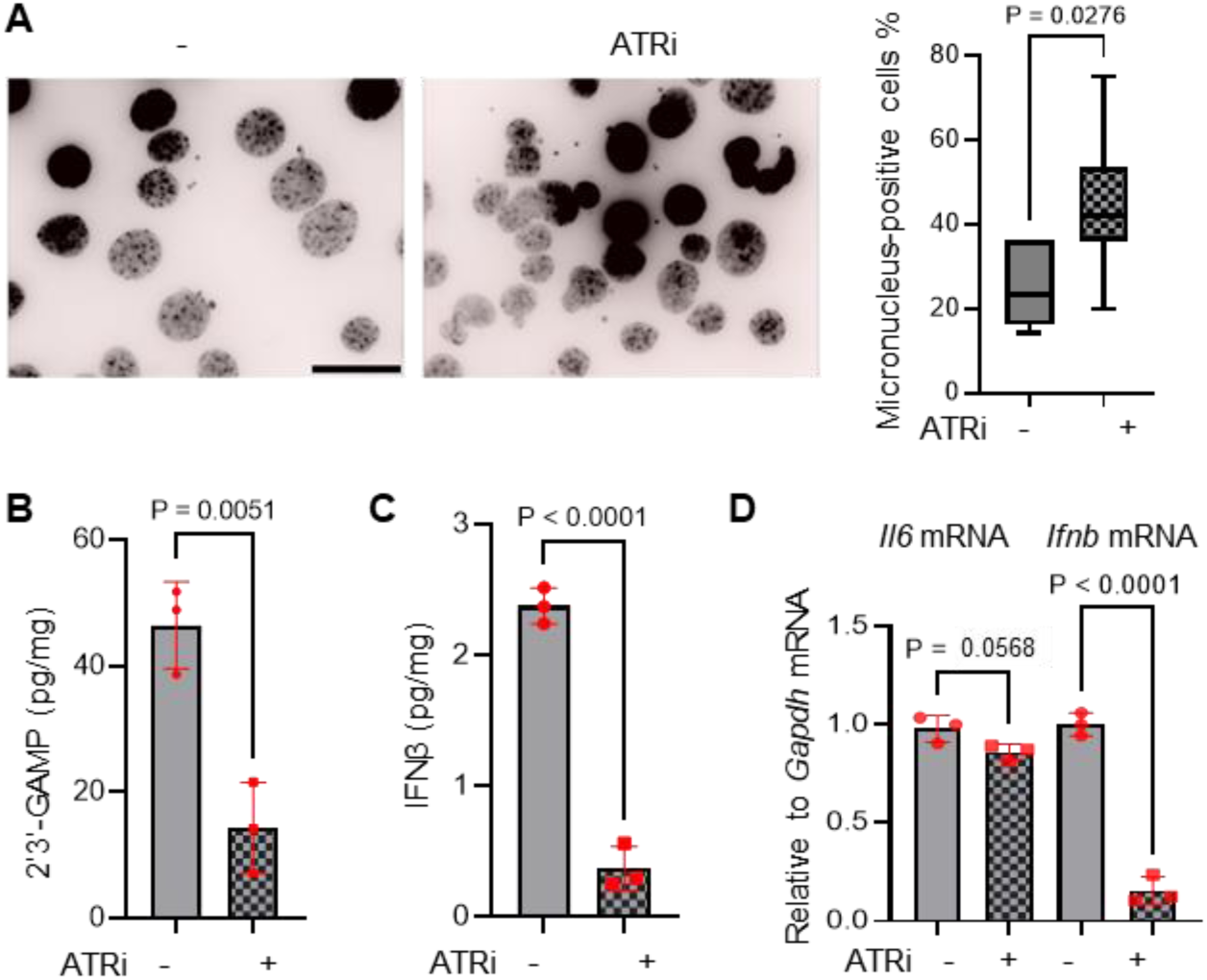
ATR signaling contributes to cGAS/STING pathway activation in TRF1-deficient MEFs. MEFs were treated with OHT, followed by the ATR inhibitor VE-821 for 24 hours. (A) Detection of micronuclei in MEFs treated with mock (ctr) or ATR inhibitor (ATRi). Cells were fixed and stained with DAPI. Scale bar, 10 μm. Quantification shows the percentage of cells with micronuclei; approximately 100 cells were counted per group. (B-C) Measurement of cellular 2’3’-cGAMP (B) and IFNβ in culture medium (C) by ELISA, with normalization to total protein concentration in the cell pellet. (D) *Il6* and *Ifnb* mRNA levels in OHT-treated MEFs exposed to mock (-) or VE-821 (ATRi), assessed by RT-qPCR. P values were calculated using Student’s t-test (A-C) and one-way ANOVA (D). Data are presented as mean ± SD from three independent experiments for panels B and C, as representative data from one of three independent biological experiments for D.

### TRF1 depletion leads to the sustained activation of NFκB/IL6 via ATM and cGAS

Nuclear DNA damage-induced ATM activation can initiate a non-canonical STING pathway, predominantly after K63-linked ubiquitin chains form on STING [32] . This modification primarily activates NFκB and drives the expression of downstream genes such as IL6, rather than triggering the typical IRF3 activation observed in cGAS-mediated DNA sensing [32]. Since ATM activation occurs subsequent to ATR activation following TRF1 ablation (Figure 1A), we investigated STING ubiquitination in TRF1-deficient MEFs. We performed immunoprecipitation (IP) for K63-linked ubiquitin in both mock- and OHT-treated MEFs, and assessed the levels of K63-ubiquitinated STING in the IP samples. Western blot analysis of input samples revealed a significant decrease in STING levels in MEFs beginning 72 hours after OHT treatment (Figure 4A, left panel). This decline is likely due to the degradation of activated STING via the proteasome and autophagy pathways [40, 41], as evidenced by the restoration of STING levels upon treatment with the proteasome inhibitor MG132 and the lysosome inhibitor Baf A1 (Figure S3).

**Figure 4:**
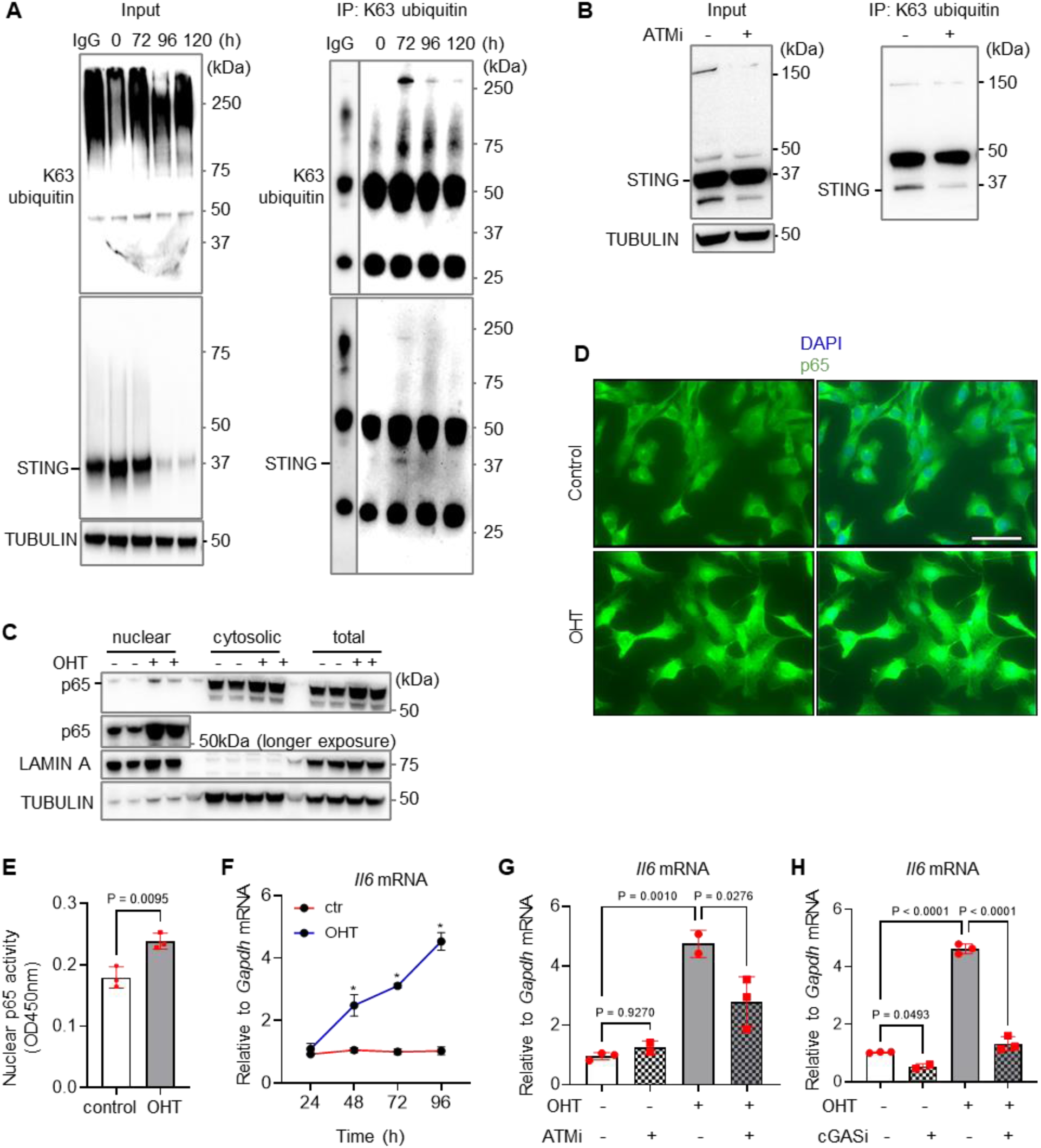
TRF1 depletion leads to STING ubiquitination and activates the NFκB pathway via cGAS and ATM. (A) MEFs were treated with OHT for the indicated times. Immunoprecipitation (IP) was performed on cell lysates using K63-linked polyubiquitin-specific antibody-conjugated Dynabeads. Input (left panel) and IP (right panel) samples were analyzed by Western blot with the indicated antibodies. IgG was used as a control (with limited exposure). Black dashes on the left indicate STING protein. (B) STING knockdown MEFs were treated with OHT, followed by the ATM inhibitor KU-55933 for 24 hours. IP was performed as described in (A). Input (left panel) and IP (right panel) samples were analyzed by Western blot. Black dashes on the left indicate STING protein. (C) Western blot analysis of p65 levels in nuclear and cytosolic fractions and whole-cell lysates from MEFs treated with mock (-) or OHT (+). A longer exposure of p65 is shown in the lower panel. Black dashes on the left indicate the protein of interest. (D) Immunofluorescence analysis of p65 (green) showing nuclear localization in OHT-treated MEFs. Nuclei were counterstained with DAPI (blue). Scale bar, 40 μm. (E) Transcriptional DNA binding activity of nuclear p65 measured in nuclear extracts from MEFs treated with mock (-) or OHT (+). (F-H) *Il6* mRNA levels were assessed by RT-qPCR: (F) in MEFs treated with mock (ctr) or OHT for the indicated times; (G) in MEFs treated with OHT for 24 hours, followed by ATM inhibitor KU-55933 (ATMi) for 72 hours; (H) in MEFs treated with OHT for 24 hours, followed by cGAS inhibitor (cGASi) for 48 hours. P values were calculated using Student’s t-test (E), one-way ANOVA (G, H), and two-way ANOVA (F) for comparisons. Data are presented as mean ± SD from three independent experiments for (E) and from a representative experiment of three independent experiments (F–H).

Notably, Western blot analysis of the K63-linked ubiquitin IP samples showed an increase in K63-linked ubiquitination of STING at 72 hours, before its degradation (Figure 4A, right panel).

To further investigate the role of ATM in STING ubiquitin modification, we treated OHT-exposed MEFs with an ATM inhibitor and performed K63-linked ubiquitin IP to assess the impact on STING ubiquitination. The efficacy of KU-55933 in inhibiting ATM was confirmed by reduced ATM phosphorylation (Figure S2B). Treatment with the ATM inhibitor significantly decreased K63-linked ubiquitination of STING in OHT-treated MEFs (Figure 4B), indicating that the ATM pathway contributes to STING ubiquitination in response to telomere dysfunction in TRF1-deficient cells.

Next, we examined the relationship between the ATM pathway and NFκB activation in TRF1-deficient cells. Immunofluorescence analysis revealed an increase in nuclear localization of p65 in MEFs after OHT treatment (Figure 4D). Consistent with this observation, quantification of nuclear p65 activity demonstrated a marked enhancement upon OHT exposure (Figure 4E). In addition, RT-qPCR analysis showed a progressive upregulation of IL6, a canonical NFκB target gene, during the course of OHT treatment (Figure 4F). Collectively, these results indicate that TRF1 depletion promotes nuclear translocation of p65 and activation of the NFκB pathway. To assess the contribution of ATM to NFκB activation, OHT-treated MEFs were exposed to the ATM inhibitor KU-55933. RT-qPCR analysis revealed a significant increase in *Il6* mRNA expression following OHT treatment, which was attenuated by KU-55933 (Figure 4G). In contrast, KU-55933 had no significant effect on the levels of *Ifnb* or *Cxcl10* mRNAs (Figure S2C).

Since the STING/TBK1 complex within the cGAS/STING pathway also contributes to NFκB activation [32], we further assessed the role of cGAS/STING signaling. Mock- or OHT-treated MEFs were treated with the cGAS inhibitor RU.521. RT-qPCR analysis showed that OHT treatment significantly upregulated *Il6* expression, which was suppressed by RU.521 (Figure 4H). Thus, in addition to ATM, cGAS also participates in NFκB signaling in TRF1-deficient cells.

### TRF1 depletion results in the activation of NFκB and IRF3 via STING

To determine whether the activation of NFκB and IRF3 is dependent on STING, we inhibited STING using the inhibitor H-151 [42, 43], and also performed STING knockdown in MEFs using siRNA and shRNA. Both western blot and RT-qPCR analyses confirmed a significant reduction in STING protein and *Sting* mRNA levels in STING-knockdown MEFs (Figure 5A-B, Figure S4A-B). Compared to non-targeting (NT) knockdown or mock controls, STING suppression, either by shRNA or H-151, resulted in decreased OHT-induced levels of p-STING, p-TBK1, p-IRF3, and p-p65 (Figure 5C-D) as assessed by western blot analysis. In turn, the OHT-induced upregulation of NFκB and IRF3/IFNβ downstream target mRNAs, specifically *Il6* and *Cxcl10*, was also attenuated by STING knockdown (siRNA or shRNA) (Figure S4C-D) and by H-151 treatment (Figure S4E). Cytokine/chemokine array analysis further supported these findings, showing that OHT-induced secretion of IFNβ and IL6 was diminished by H-151 treatment (Figure 5E,F). Notably, treatment with H-151 also reduced the levels of other cytokines, including TNF-α, IL-1β, and IFN-γ, in OHT-treated MEFs (Figure S4F). Collectively, these results indicate that activation of NFκB and IRF3 is critically dependent on STING signaling. Suppression of STING significantly reduces the activation of key NFκB and IRF3 pathway proteins and downstream gene expression of IL6 and IFNβ. Furthermore, our data indicate that STING broadly modulates the inflammatory response, as indicated by the reduction of additional cytokines such as TNF-α, IL-1β, and IFN-γ upon STING inhibition

**Figure 5:**
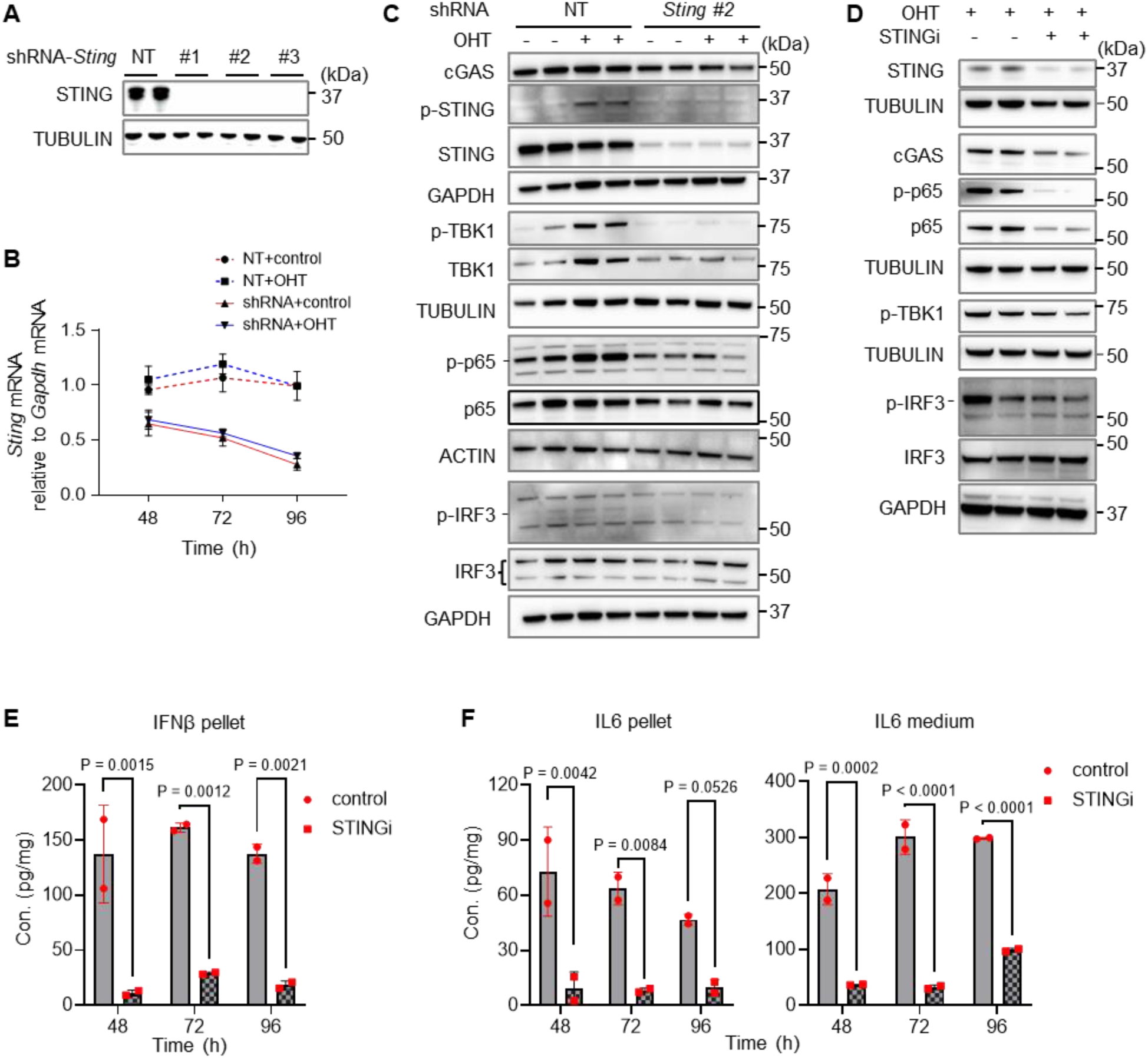
Activation of the NFκB/IL6 and IRF3/IFNβ pathways is STING-dependent in TRF1-deficient MEFs. (A) Western blot analysis confirming STING knockdown. MEFs were transfected with either non-targeting shRNA control (shNT) or *Sting* shRNA lentivirus for 72 hours. (B) shRNA-mediated STING knockdown MEFs were treated with mock or OHT for the indicated times, and knockdown efficiency was assessed by RT-qPCR. (C) Whole cell lysates harvested at 96 hours were analyzed by Western blot with the indicated antibodies. Black dashes on the left denote the position of the protein of interest. (D) MEFs were treated with OHT for 96 hours in the presence (+) or absence (–) of the STING inhibitor H-151. H-151 was administered 24 hours post-initiation of OHT treatment. Cell lysates were analyzed by Western blot with the indicated antibodies. (E–F) IFNβ and IL6 protein levels were measured by cytokine array in both cell pellets and culture medium from OHT-treated MEFs for indicated times. H-151 was administered 24 hours post-initiation of OHT treatment. Values were normalized to total protein concentration in the cell pellet. P values were calculated using two-way ANOVA for multiple comparisons. Data are presented as mean ± SD from a representative experiment out of three independent experiments.

### ATM/STING & ATR/cGAS/STING axis synergistically regulate the NFκB pathway in TRF1-deficient cells

Thus far, we have shown that both the ATM/STING and cGAS/STING pathways contribute to NFκB-mediated inflammatory responses upon TRF1 depletion. To further explore their interplay, we investigated how these pathways converge to regulate NFκB activation in NFκB-mediated inflammatory responses in TRF1-deficient MEFs. TRF1-deficient MEFs were treated with the ATM inhibitor KU-55933 and/or the cGAS inhibitor RU.521, followed by measurement of *Il6* mRNA expression via RT-qPCR. Both inhibitors individually reduced *Il6* expression, and their combined application resulted in an even greater reduction in *Il6* mRNA levels (Figure 6A).

**Figure 6.**
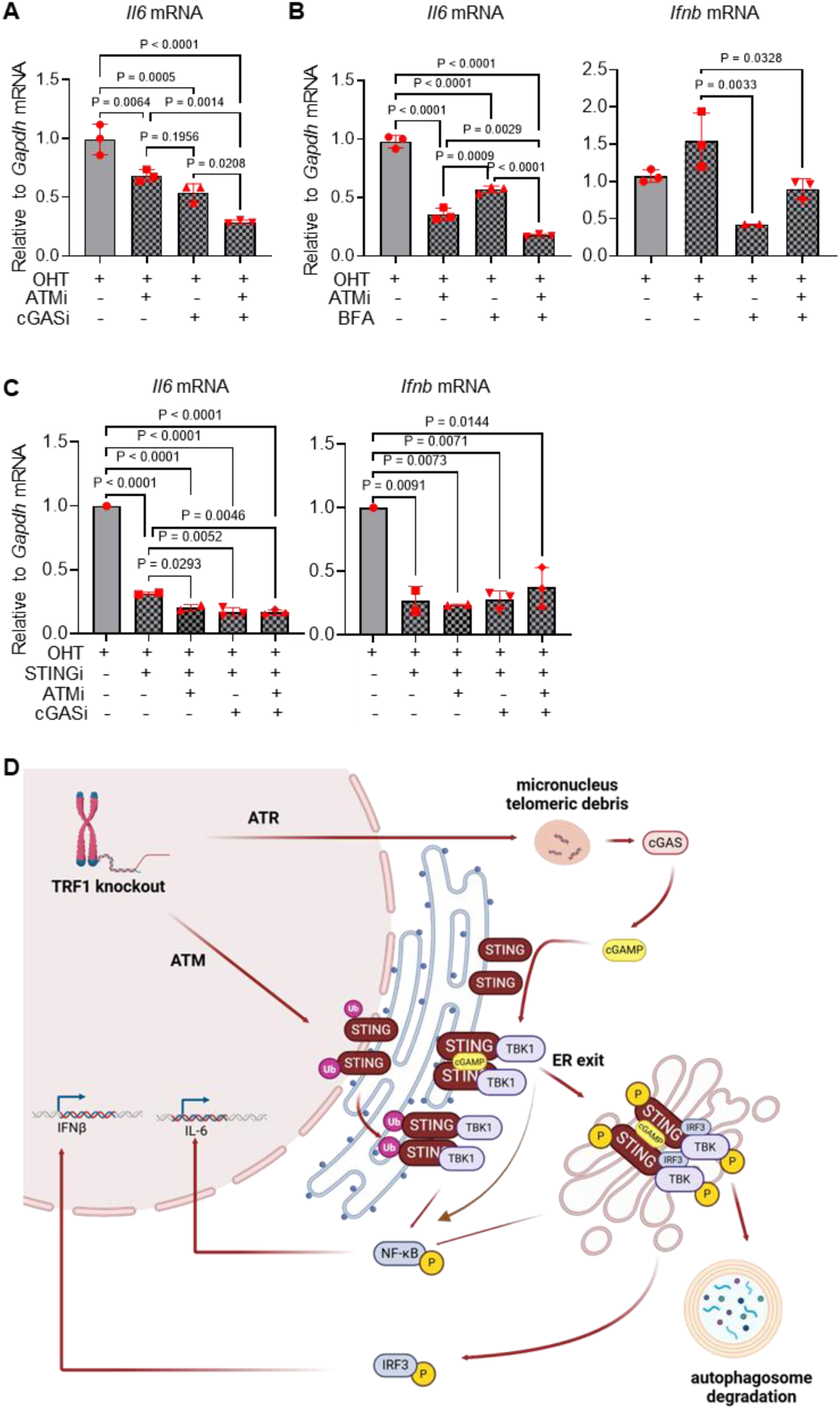
ATM and cGAS synergistically activate STING downstream pathway genes in TRF1-deficient MEFs. (A) MEFs were treated with OHT to induce TRF1 deletion, followed by treatment with ATM inhibitor KU-55933, cGAS inhibitor RU.521, or both for 72 hours. *Il6* mRNA expression was assessed by RT-qPCR. (B) MEFs were treated with OHT, KU-55933 (ATMi, 72 hours), and/or BFA (ER exit inhibitor, 3 hours). *Il6* and *Ifnb* mRNA levels were measured by RT-qPCR. (C) MEFs were treated with OHT to induce TRF1 deletion, followed by the STING inhibitor H-151 alone or in combination with KU-55933 (ATMi) and/or RU.521 (cGASi) for 72 hours. *Il6* and *Ifnb* mRNA expression was measured by RT-qPCR. (D) Schematic illustration of the proposed mechanism for ATM/STING and ATR/cGAS/STING pathway activation following TRF1 depletion. P values were calculated using one-way ANOVA for multiple comparisons. Data are presented as mean ± SD from a representative experiment out of three independent experiments (A–C).

IRF3 activation by the cGAS/STING pathway predominantly occurs in the Golgi after STING exits the ER, while STING-dependent NFκB activation may begin as early as in the ER and continue in the Golgi after ER exit [22, 44, 45]. Thus, ATM- and cGAS-mediated STING activation may cooperatively stimulate the expression of downstream genes regulated by NFκB and/or IRF3 through both the ER and post-ER pathways. To test this, we treated OHT-induced MEFs with Brefeldin A (BFA), an inhibitor of ER exit, and/or the ATM inhibitor, KU-55933, then assessed NFκB and IRF3 target gene expression. KU-55933 markedly reduced *Il6* levels, BFA produced a moderate decrease, and combined treatment led to a synergistic reduction in *Il6* mRNA levels (Figure 6B). These results suggest that ATM primarily activates NFκB signaling in the ER, but also collaborates with STING after ER exit to modulate NFκB activity. In contrast, KU-55933 had little effect on *Ifnb* mRNA levels, whereas BFA treatment significantly reduced its levels (Figure 6B), indicating that ER exit is essential for IRF3 downstream gene activation.

To further dissect the relationships among the ATM, cGAS, and STING pathways, TRF1-deficient MEFs were pre-treated with the STING inhibitor H-151 prior to treatment with ATM and/or cGAS inhibitors. RT-qPCR analysis revealed that STING inhibition significantly decreased the levels of both *Il6* and *Ifnb* mRNAs. While the addition of ATM and/or cGAS inhibitors caused a slight further decrease in the levels of *Il6* and *Ifnb* mRNAs, these remained largely unaffected by the additional treatments (Figure 6C). These findings underscore the critical role of STING in mediating the induction of IL6 and IFNβ by ATM and cGAS signaling.

Collectively, our data suggest that in response to telomere dysfunction in TRF1-deficient cells, the ATM pathway may promote STING ubiquitination and subsequent NFκB activation in the ER. Following ATR activation and micronuclei rupture, the cGAS pathway senses cytosolic telomeric/DNA fragments, leading to STING activation and its exit from the ER, which in turn activates IRF3 and NFκB signaling. These two STING-dependent mechanisms function synergistically to modulate inflammatory responses in TRF1-deficient MEFs (Figure 6D).

## Discussion

Replication stress activates a DDR and serves as a fundamental driver of cellular senescence and aging [46]. Telomeres, characterized by their repetitive guanine-rich sequences and propensity to form alternative DNA structures, are particularly vulnerable to replication stress and DNA damage [11]. The key telomeric protein TRF1 plays a pivotal role in facilitating proper telomere replication and protecting chromosome ends. TRF1 deficiency increases telomere replication stress and fragility, leading to DNA damage, activation of the DNA damage response (DDR), and ultimately cellular senescence [10, 11]. In mice, TRF1 loss induces a persistent inflammatory response, which drives premature tissue degeneration and promotes the development of lung and kidney fibrosis [10, 47, 48]. However, the molecular mechanisms linking telomere replication stress and dysfunction to inflammation require further investigation. Here, we demonstrate that telomere replication stress and dysfunction caused by TRF1 depletion can elicit a STING-dependent inflammatory response, mediated through ATR- and ATM-driven DNA damage response pathways.

The canonical cGAS/STING pathway is a critical component of the innate immune response to DNA damage [14], initiated by sensing exposed double-stranded DNA in the cytosol [49]. During this process, ATR promotes the rupture of micronuclei, leading to the release of cytosolic DNA and subsequent activation of the cGAS/STING pathway [15, 39, 41]. Beyond its role in inflammation through downstream signaling, cGAS/STING activation also facilitates the clearance of cytosolic DNA and viral particles [41]. Notably, telomeres are recognized as hotspots for innate immune activation and inflammation [50]. While micronuclei generated by general genome replication stress or chromosome mis-segregation do not typically activate the cGAS/STING pathway [49], those arising from telomeric DNA double-strand breaks or telomere uncapping can indeed trigger this pathway [50–52]. Furthermore, premature senescence induced by telomeric DNA double-strand breaks, such as those created by CRISPR/Cas9, is driven by cGAS/STING activation, independent of telomere shortening [51]. Here, we show that TRF1 depletion results in the accumulation of cytosolic and micronuclear telomeric fragments, increased levels of cytosolic cGAS, 2’3’-cGAMP, phosphorylated STING, phosphorylated IRF3 and its nuclear translocation, as well as elevated IFNβ expression in MEFs. In contrast, treatment with an ATR inhibitor prevents micronuclear rupture, as indicated by an increased numbers of micronuclei and reduced levels of 2’3’-cGAMP, produced in response to cytosolic dsDNA, in ATR inhibitor–treated TRF1-deficient cells compared to mock-treated controls.

Additionally, ATR inhibition lowers IFNβ levels more substantially than IL6 in TRF1-deficient cells. These findings suggest that ATR activation promotes micronuclear rupture in TRF1-deficient cells, thereby initiating cGAS/STING signaling, which predominantly enhances IRF3/IFNβ pathway activity. Consistent with the findings of Takaki et al. [49], we propose that the DNA fragments present in micronuclei are newly synthesized, unprotected DNA generated as a consequence of replication stress from TRF1 depletion, which in turn activates the cGAS/STING pathway.

Previous studies have shown that TRF1 deletion activates either ATR or ATM signaling pathway [10, 11]. Building on this, our results indicate that ATR activation occurs prior to ATM activation following TRF1 depletion. We observed a significant increase in micronuclei formation and 53BP1 foci in TRF1-deficient cells, suggesting the presence of chromosome fragments and DNA double-strand breaks. This implies that replication stress induced by TRF1 loss disrupts telomere and genomic stability, subsequently activating the ATM pathway.

ATM activation in response to DNA damage is known to promote inflammation, primarily through the NFκB pathway and non-canonical STING activation via K63-linked ubiquitination [30–32]. Consistent with this regulatory paradigm, TRF1-deficient cells exhibited higher levels of K63-linked ubiquitin-modified STING compared to TRF1-proficient cells. Additionally, these cells displayed hallmarks of NFκB activation, including increased nuclear translocation and activity of p65, as well as increased transcription of the NFκB target gene *Il6*. Notably, while the expression levels of the IRF3 target *Ifnb* mRNA showed a rapid but transient increase, IL6 activation was both robust and sustained in TRF1-deficient MEFs. Even as STING levels began to decline after OHT treatment, IL6 levels remained relatively high. These findings support the hypothesis that the ATM/STING/NFκB/IL6 axis drives chronic inflammation in response to telomere dysfunction via STING ubiquitination. This possibility is further substantiated by the observation that both IL6 expression and STING K63-linked ubiquitination in TRF1-deficient cells are reduced by ATM inhibition. However, the cGAS/STING pathway also appears to contribute to NFκB activation, as IL6 production was suppressed by a cGAS inhibitor. Moreover, combined treatment with ATM and cGAS inhibitors synergistically reduced IL6 levels in TRF1-deficient cells. Together, these results suggest that the ATM/STING and ATR/cGAS/STING pathways cooperate to modulate inflammatory responses in TRF1-deficient MEFs.

Our data demonstrate that STING plays a pivotal role in mediating the inflammatory response triggered by TRF1 depletion. We have shown that OHT-induced increases in the phosphorylation of STING, TBK1, IRF3, and p65, as well as their downstream targets IL6 and IFNβ were significantly attenuated by either STING knockdown or pharmacological inhibition. Furthermore, inhibition of ATM and/or cGAS failed to markedly reduce IL6 and IFNβ expression when STING was inhibited beforehand. These findings indicate that STING acts as a key regulator, integrating signals from both the ATM and cGAS pathways in the inflammatory cascade induced by TRF1 depletion. Thus, the convergence of ATM and cGAS activation on STING amplifies the inflammatory response following TRF1 loss.

Based on these observations, we propose a model in which TRF1 depletion induces telomere replication stress, resulting in DNA damage, sequential activation of the ATR- and ATM-dependent DDR pathways, and the formation of micronuclei. ATR activation triggers micronuclei rupture, releasing cytosolic telomeric and genomic DNA fragments that activate the cGAS/STING pathway. This activation promotes STING translocation from the endoplasmic reticulum, facilitating the nuclear localization of NFκB and IRF3 and the upregulation of their respective target genes [21]. ATM activation promotes K63-linked ubiquitination of STING, which preferentially activates the NFκB pathway and sustains inflammation via the ATM/STING/NFκB/IL6 axis. Together, these two STING-dependent pathways enable efficient signal transduction in response to DNA damage and the presence of cytosolic DNA. As STING inhibition can disrupt this inflammatory response, targeting STING may hold therapeutic potential for conditions associated with telomere replication stress and dysfunction, thereby restoring normal cellular function.

## MATERIALS AND METHODS

### Cell line, cell culture, and treatment

MEFs carrying the SV40LT TRF1^F/F^ Rosa26Cre-ER^T1^ alleles were provided by Dr. Titia de Lange [11]. Cells were cultured at 37°C with 5% CO₂ in DMEM (Thermo Fisher Scientific, 11965118) supplemented with 10% fetal bovine serum (FBS; Millipore Sigma, F0926) and 1% glutamine (Thermo Fisher Scientific, 25030081). TRF1 deletion was induced by treatment with 1 μM 4-hydroxytamoxifen (OHT), which enabled Cre-mediated excision of exon 1 of the mouse TRF1 gene.

The following chemicals were used for cell treatments in the experiments: ATM inhibitor KU-55933 (Abcam, 120637, 10 μM), ATR inhibitor VE-821 (Selleck, S8007, 2.5 μM); cGAS inhibitor RU.521 (Invivogen, inh-ru521, 10 μM), STING-specific inhibitor H-151 (MedChem Express, HY-112693, 7.5 μM), STING ER exit inhibitor Brefeldin A (BFA; Thermo Fisher Scientific, 00-4506-51, 3.0 µg/mL), proteasome inhibitor MG132 (Millipore Sigma, M7449, 5 μM), lysosome inhibitor Bafilomycin A1 (Baf A1; Millipore Sigma, 19-148, 100 nM), and replication stress inducer Aphidicolin (Millipore Sigma, 178273, 0.2 μM).

### STING knockdown

STING-specific small interfering RNAs (siRNAs) (SASI_Mm02_00429135, SASI_Mm02_00429136, SASI_Mm01_00247170) and a non-targeting siRNA control (siNT) were obtained from Millipore Sigma. siRNA transfection was performed using the INTERFERIN transfection reagent (Polyplus, 101000036). Cells were harvested 2 to 3 days post-transfection, and knockdown efficiency was assessed by reverse transcription quantitative PCR (RT-qPCR) and western blot analysis.

STING small hairpin RNA (shRNA) lentiviral plasmids (Millipore Sigma, SHCLNG-TRCN0000346266, SHCLNG-TRCN0000346319, SHCLNG-TRCN0000346320), along with the packaging plasmid (pCMV Δ8.2) and envelope plasmid (pCMV VSVG), were used to generate lentivirus in 293T cells. For the transfection mixture, 500 μL of FBS-free DMEM was combined with 5 μg of shRNA plasmid, 4.5 μg of packaging plasmid, and 0.5 μg of envelope plasmid. This mixture was then diluted with 45 μL of Plus DNA In Vitro Transfection Reagent (SignaGen, SL100499) and incubated at room temperature for 15 minutes. Subsequently, 1 mL of the transfection mixture was added to a dish containing 5 mL of fresh medium. After 5 hours, the medium was replaced with fresh medium. The culture supernatant was collected 48-72 hours post-transfection. Lenti-X GoStix Plus (TAKARA, 631280) was used for rapid assessment of lentiviral titer. The supernatant was filtered through a 0.45 μm filter to remove cell debris, mixed with polybrene (Millipore Sigma, TR-1003-G) to a final concentration of 10 μg/mL, and used to infect MEFs. The next day, puromycin (Invivogen, ant-pr-1) was applied at a final concentration of 2 μg/mL in fresh medium and continued for 3 to 5 days. Knockdown efficiency of STING was confirmed by RT-qPCR and western blot analysis.

### Immunofluorescence (IF), metaphase spread preparation, and telomere fluorescent in situ hybridization (telomere-FISH)

IF was performed as previously described, with minor modifications [53]. Cells were seeded and cultured on 20 mm round coverslips. They were fixed with 4% paraformaldehyde (Thermo Fisher Scientific, J19943-K2) at 4°C, permeabilized with 0.2% Triton X-100 (Millipore Sigma, X100-100ML), and blocked with 5% FBS at room temperature for 1 hour. Cells were then incubated overnight at 4°C with primary antibodies, cGAS (Invitrogen, # PA5-121188, 1:50), STING (Cell Signaling Technology, 90947, 1:3200), p65 (Cell Signaling Technology, 8242, 1:400), p-IRF3 (Cell Signaling Technology, 29047, 1:100), or Calregulin (Santa Cruz, sc-166837, 1:50). This was followed by incubation with Alexa Fluor-labeled secondary antibodies (Abcam) for 1 hour at 37°C. For combined telomere FISH, slides were washed with 1x PBS (KD medical, RGF-3210) for 15 minutes, fixed in 2% paraformaldehyde at room temperature for 10 minutes, and then dehydrated through a series of ethanol (Millipore Sigma, 1009831000) solutions. After a brief air-drying, the slides were processed for telomere-FISH.

For metaphase spread preparation, cells were treated with 0.1 μg/mL colcemid (Millipore Sigma, 10295892001) for 2 hours at 37°C, harvested, and incubated in 75 mM KCl (Fisher Scientific, 01-911-512) for 15-20 minutes at 37°C. Cells were then fixed in ice-cold methanol (Fisher Scientific, A452-4) and glacial acetic acid (Sigma, A6283-100ML) at a 3:1 volume ratio, and dropped onto glass slides.

For telomere-FISH, metaphase spreads or cells were hybridized with either a Cy3-labeled or FITC-labeled (CCCTAA)_3_ PNA probe (0.5 μg/mL, Panagene, F1002, F1009), washed, and mounted with ProLong Gold anti-fade reagent (Invitrogen, P36962), as previously described [53]. Images were acquired using a fluorescence microscope (Axio2; Carl Zeiss) equipped with Cytovision software (Applied Imaging).

### RNA isolation and RT-qPCR analysis

Total RNA was extracted with Trizol reagent (Invitrogen, 15596018) in the presence of an RNase inhibitor, as previously described [54]. Briefly, samples were mixed with 1 mL of Trizol and 200 μL of chloroform (Millipore Sigma, C2432-500ML), then centrifuged at 12,000 g for 15 minutes at 4°C. The aqueous phase was collected and RNA was precipitated by adding an equal volume of isopropanol (Thermo Fisher Scientific, T036181000CS). RNA pellets were collected by centrifugation at 12,000 g for 10 minutes, washed with 75% ethanol, air-dried, and resuspended in RNase-free water. Reverse transcription was performed using PrimeScript™ RT Master Mix (TAKARA, RR036A), and qPCR was conducted with iTaq Universal SYBR Green supermix (Bio-Rad, 1725121). Relative mRNA expression was calculated using the 2^-ΔΔCT^ method and normalized to *Gapdh* mRNA. Primer sequences used for RT-qPCR are listed in Table S1.

### Cell fractionation

Nuclear and cytosolic fractionation was performed as previously described [55]. Approximately 10 million cells were transferred to a 15 mL tube and washed twice with 1x PBS. After centrifugation at 21,000 g for 10 seconds, the supernatant was discarded, and the cell pellet was resuspended by pipetting five times in 1 mL of ice-cold 1x PBS lysis buffer containing 0.1% NP-40 (Santa Cruz, sc-281108). Cell lysate was centrifuged at 21,000 g for 10 seconds to pellet the nuclei. Half of the resulting supernatant was collected as the “cytosolic fraction”, while the nuclear pellet was washed with 1 mL of lysis buffer. The nuclear pellet was then resuspended in 1× Laemmli buffer (Bio-Rad, 1610747) and designated as the “nuclear fraction”. The cytosolic fraction was mixed with 4× Laemmli buffer. All protein samples were boiled at 95 °C for 10 minutes. Western blot analysis was performed using equal amounts of protein from the nuclear and cytosolic fractions.

### Western blot analysis and antibodies

Cells were resuspended in a buffer containing 0.1% SDS (Fisher Scientific, MT-46040CI), 0.05 M Tris-HCl (pH 8) (KD Medical, RGF-3360), and 0.01 M EDTA (pH 8) (Quality Biological, 351-027-101). The samples were sonicated in 30-second pulses for 10 cycles after being pipetted several times. Following sonication, samples were mixed with 4× Laemmli buffer containing 50 mM DTT (Bio-Rad, 1610610) and boiled at 95 °C for 10 minutes. Proteins were separated by molecular weight using Bis-Tris gels (Invitrogen, NuPAGE) and transferred to PVDF membranes (Bio-Rad, 1620177). Membranes were incubated with primary antibodies, followed by secondary antibodies, and visualized using the ChemiDoc imaging system (Bio-Rad). For sequential probing, membranes were incubated with western blot stripping buffer (Thermo Scientific, 46430) before re-incubation with additional primary antibodies targeting proteins of similar molecular weight.

The following primary antibodies were utilized for western blot, including p-IRF3 (Cell Signaling Technology, 4947, 1:1000), IRF3 (Cell Signaling Technology, 4302, 1:1000), TRF1 (Abcam, 192629, 1:1000), p-TBK1 (Cell Signaling Technology, 5483, 1:1000), TBK1 (Cell Signaling Technology, 3504, 1:1000), p-STING (Cell Signaling Technology, 72971, 1:1000), STING (Cell Signaling Technology, 50494, 1:1000), p-CHK1 (Cell Signaling Technology, 2348, 1:1000), CHK1 (Cell Signaling Technology, 2360, 1:1000), p-CHK2 (Bioss, 3721, 1:1000), CHK2 (BD, 611570, 1:500), p-ATM (Santa Cruz, 47739, 1:500), ATM (Cell Signaling Technology, 2873, 1:1000), p-ATR (Cell Signaling Technology, 2853, 1:1000), ATR (Abcam, 2905, 1:1000), p-p65 (Cell Signaling Technology, 3033, 1:1000), p65 (Cell Signaling Technology, 8242, 1:1000), cGAS(Cell Signaling Technology, 31659, 1:1000), LAMIN A (Abcam, 26300, 1:1000), β-TUBULIN (Abcam, 6046, 1:500), and GAPDH (Abclonal, AC027, 1:2000).

### Immunoprecipitation (IP)

STING, ubiquitin, and TRAF6 plasmids were obtained from Addgene (#102598, #18712, and #21624). MEFs were transfected with the plasmids 24 hours prior to cell collection using the jetPRIME transfection reagent (Polyplus, 101000027). Cells were seeded in 10-cm dishes, and for each dish, 250 μL of jetPRIME buffer was mixed with 2 μg of STING plasmid, 2 μg of ubiquitin plasmid, 1 μg of TRAF6 plasmid, and 10 μL of jetPRIME reagent. The mixture was incubated at room temperature for 10 minutes before use. Prior to transfection, 5 mL of the old medium was removed from each dish.

Cell lysate was incubated with IP lysis buffer (Thermo Fisher Scientific, 87787) supplemented with 10 mM iodoacetamide (SIGMA, I1149-5G) and protease and phosphatase inhibitors (Thermo Fisher Scientific, 78442). Supernatant was collected by centrifugation at 8000g for 10 minutes. For immunoprecipitation, approximately 50 μL of protein G or protein A Dynabeads (Invitrogen, Thermo Fisher Scientific, 10003D and 10008D) were incubated with either K63-linked specific polyubiquitin antibody (Cell Signaling Technology, 5621) or STING antibody (Cell Signaling Technology, 50494) at 4°C for 1 hour with rotation to allow antibody binding. Next, 200 μL of supernatant was added to the Dynabeads-antibody mixture and incubated with rotation. After washing, the Dynabeads-antibody-protein complexes were resuspended in 1× Laemmli buffer and designated as IP samples. The remaining supernatant was reserved as “input” and mixed with 4× Laemmli buffer. Both IP and input samples were boiled at 70°C for 10 minutes and subsequently analyzed by western blot.

### Nuclear NFκB p65 activity

Nuclear p65 activity was measured using the NFκB p65 Transcription Factor Assay Kit (Abcam, 133112). Briefly, nuclei were isolated with a nuclear extraction kit (Abcam, ab113474) and centrifuged at 14,000 g for 10 minutes at 4°C. The supernatant was collected and diluted to a concentration of 0.5 mg/mL. Diluted samples were incubated overnight at 4°C with the Transcription Factor Binding Assay Buffer. After washing, the p65 primary antibody was added and incubated at room temperature for 1 hour. Following incubation with a secondary antibody and additional washes, the developing solution was added for 30 minutes. The reaction was stopped with stop solution, and p65 activity was quantified by measuring absorbance at 450 nm (OD450).

### Cellular cGAMP measurement

Cellular cGAMP level was measured using the 2’3’-cGAMP ELISA Kit (Cayman Chemical, 76292-890) according to the manufacturer’s instructions. Briefly, cells were lysed in 400 μL of mammalian protein extraction reagent (Thermo Fisher Scientific, 78503). The lysates were vortexed at 850 rpm for 5 minutes, followed by ultrasonic disruption for 10 minutes. After centrifugation at 14,000 g for 10 minutes, the supernatant was collected and adjusted to 2 mg/mL with 1× buffer C. For the ELISA, 50 μL of sample was combined with 50 μL cGAMP tracer and 50 μL antiserum in each well of a mouse anti-rabbit IgG-coated plate, and incubated overnight at 4°C. After washing, 175 μL of TMB substrate was added and incubated for 30 minutes at room temperature. The reaction was stopped with 75 μL of stop solution, and absorbance was measured at 450 nm (OD450). Standard curves were generated using a four-parameter logistic fit to calculate cGAMP concentrations in each sample.

### Cytokine/Chemokine measurement

Cells were seeded in 6-well plates, and both the cells and culture medium were collected. Cells were lysed using RIPA buffer (Thermo Fisher Scientific, 89901) supplemented with protease and phosphatase inhibitors. After centrifugation at 14,000 rpm for 15 minutes at 4°C, the supernatant was collected. Both cell lysates and medium samples were analyzed for cytokine and chemokine levels by Eve Technologies.

IFNβ levels in the cell culture medium were measured using the Mouse IFN-Beta ELISA Kit (PBL Assay Science, 42410-1). Briefly, culture medium was collected in a 15 mL tube and centrifuged at 2,000 rpm for 5 minutes to remove cell debris; the resulting supernatant was used for the assay. In each well, 50 μL of serum buffer and 50 μL of standard or sample were added and incubated for 1 hour at room temperature with shaking at 650 rpm. After washing, 50 μL of antibody solution and 50 μL of HRP solution were added sequentially and incubated for 30 minutes and 10 minutes, respectively. Following another wash, 100 μL of TMB substrate was added and incubated for 10 minutes in the dark. The reaction was stopped with 100 μL of stop solution, and absorbance was measured at 450 nm (OD450). Results were normalized to the protein concentration of the corresponding cell lysate.

### Statistical analysis

Statistical analysis was performed using Student’s t-test for comparisons between two groups, and ANOVA for comparisons among multiple groups. P values less than 0.05 were considered statistically significant. All analyses were conducted using GraphPad Prism 10 (GraphPad Software, Inc.). Data are presented as mean ± standard deviation (SD).

### Data availability

The authors declare that data supporting the findings of this study are available within the paper and its supplementary information files. Requests for materials should be made to Dr. Yie Liu.

## AUTHOR CONTRIBUTIONS

Y.L. M.G. and W.Z. conceived the project. Y.L., W.Z., and Y.W. designed the experiments. W.Z., Y.W., and Y.G. performed experiments, statistical analysis, and/or data analysis; Y.L. and M.G. provided project administration, resource, and supervision support. Y.L., M.G., and W.Z. wrote the manuscript, and all authors read, reviewed, and/or edited the manuscript.

## ACKNOWLEDGMENTS

We sincerely thank Dr. Titia de Lange for providing SV40LT TRF1^F/F^ Rosa26Cre-ER^T1^ mouse embryonic fibroblasts. This research was supported [in part] by the Intramural Research Program of the National Institutes of Health (NIH). The contributions of the NIH author(s) were made as part of their official duties as NIH federal employees, are in compliance with agency policy requirements, and are considered Works of the United States Government. However, the findings and conclusions presented in this paper are those of the author(s) and do not necessarily reflect the views of the NIH or the U.S. Department of Health and Human Services.

## SUPPLEMENTARY FIGURES

**Figure S1.**
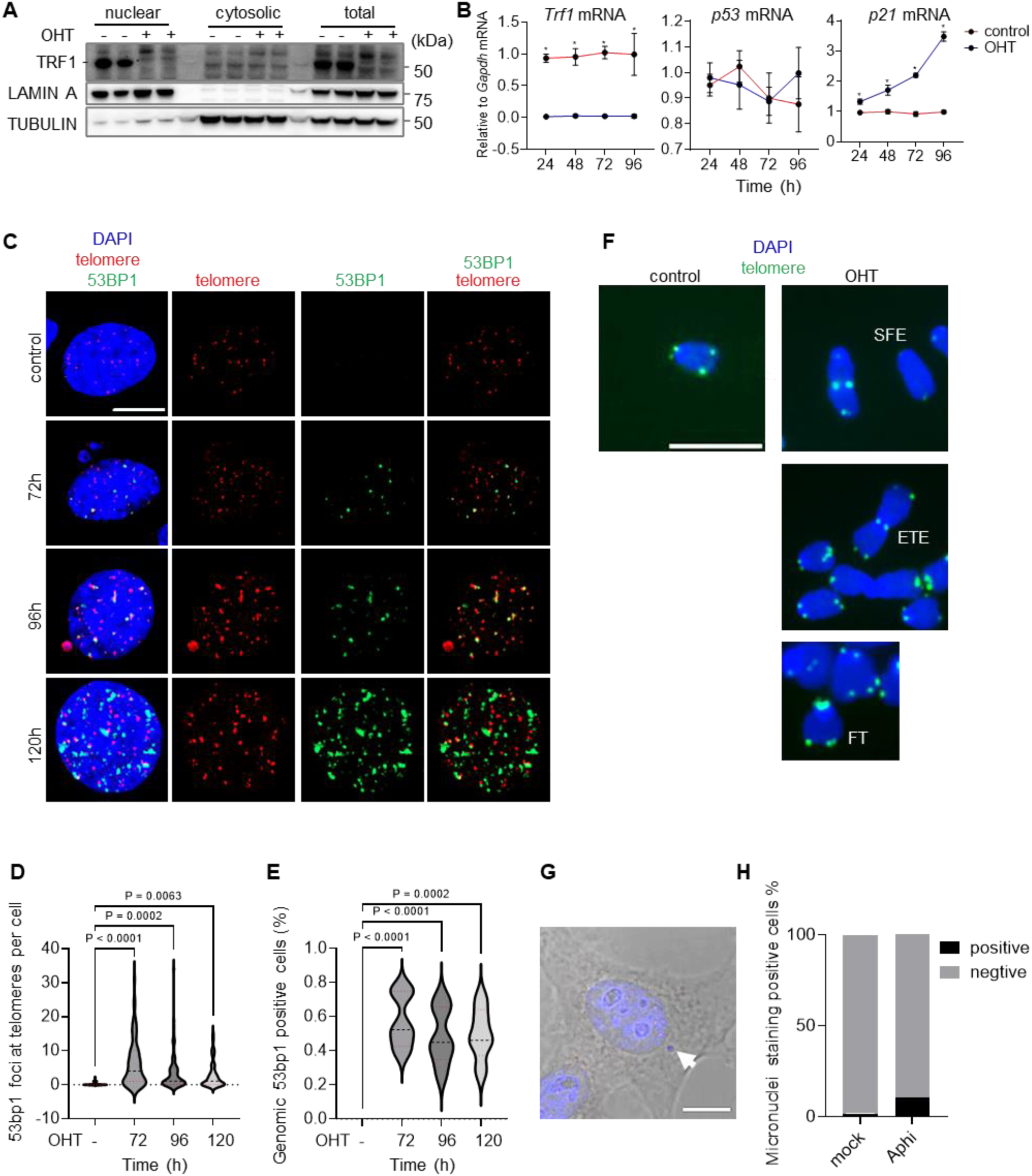
TRF1 depletion triggers a DNA damage response in MEFs. (A) MEFs were treated with OHT for 96 hours. TRF1 protein levels were analyzed by Western blot in nuclear and cytoplasmic fractions and whole cell lysates. Black dashes on the left indicate the position of the protein of interest. (B) *Trf1*, *p53*, and *p21* mRNA levels were measured by RT-qPCR in MEFs treated with OHT for indicated times. (C) 53BP1 foci at telomeres and throughout the genome were detected in OHT-treated MEFs using immunofluorescence-telomere FISH with a telC-Cy3 probe. Representative images show 53BP1 (green), telomeric DNA (red), and total DNA (DAPI, blue). Scale bar, 10 μm. (D) Quantification of 53BP1 foci at telomeres per cell. (E) Percentage of cells positive for genomic 53BP1 foci. (F) Telomere abnormalities, including telomere signal-free ends (SFE), fragile telomeres (FT), and end-to-end fusions (ETE), were detected in metaphase spreads of OHT-treated MEFs by telomere FISH using the telC-FITC probe (green). DNA was counterstained with DAPI (blue). Scale bar, 4 μm. (G-H) OHT-treated MEFs were exposed with a low dosage of aphidicolin for 24 hours. (G) Representative telomere FISH image showing a cytosolic telomere signal (arrow) using the telC-Cy3 probe. Scale bar, 10 μm. (H) Quantification of the percentage of cells positive for micronuclei. P values were calculated using one-way ANOVA (D, E) or two-way ANOVA (B) for multiple comparisons. Data are presented as mean ± SD from a representative experiment out of three independent experiments (B, * denotes P values <0.05), or from approximately 100 cells per group (D, E).

**Figure S2.**
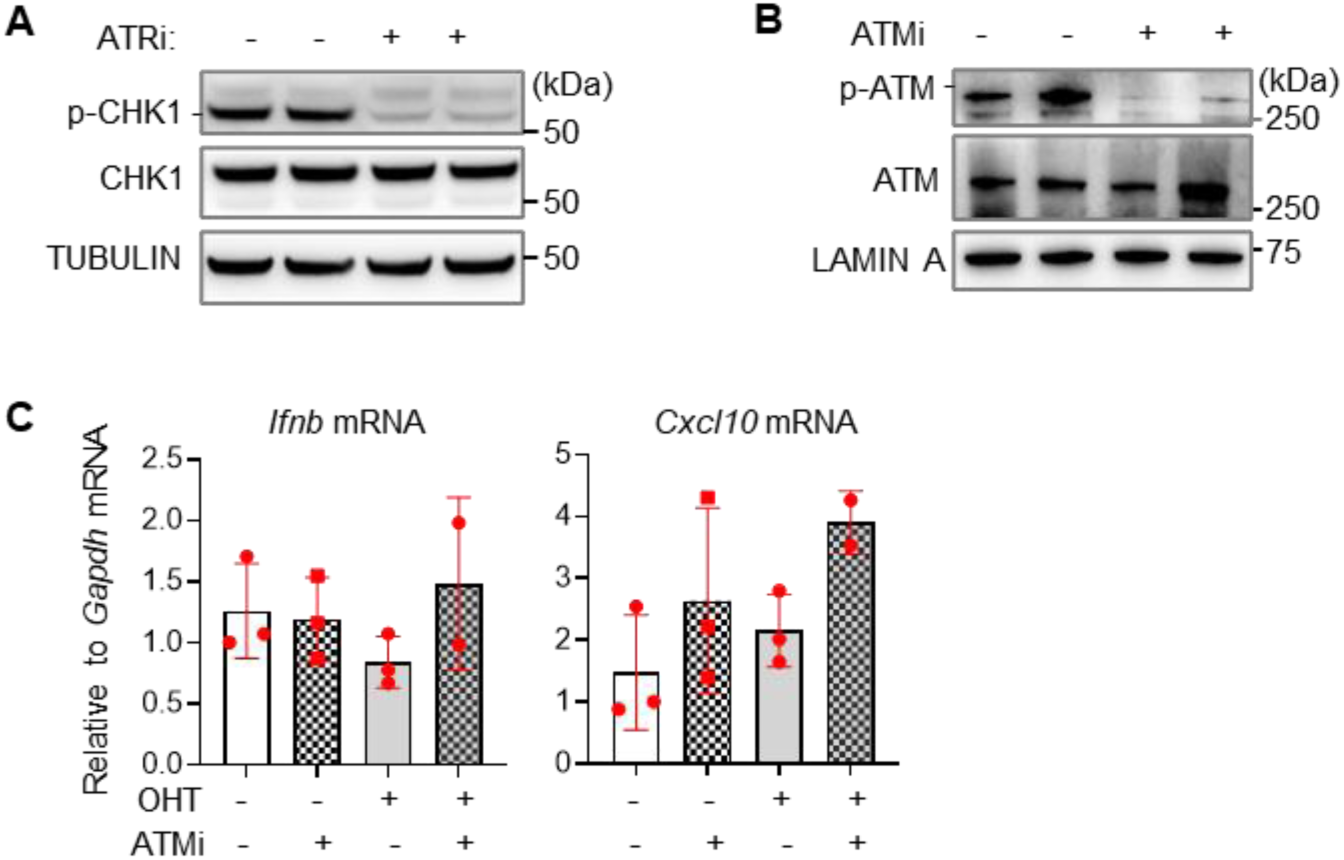
ATR and ATM pathway inhibition in TRF1-deficient MEFs and its effect on inflammatory gene expression. (A) Western blot analysis demonstrating the efficacy of ATR pathway inhibition in TRF1-deficient MEFs treated with the ATR inhibitor VE-821 for 4 hours. (B) Western blot analysis demonstrating the efficacy of ATM pathway inhibition in MEFs treated with the ATM inhibitor KU-55933 for 1 hour. Black dashes on the left indicate the position of the protein of interest. (C) RT-qPCR analysis of *Ifnb* and *Cxcl10* mRNA expression in TRF1-deficient MEFs treated with the ATM inhibitor KU-55933. Data are presented as mean ± SD from a representative experiment out of three independent experiments.

**Figure S3.**
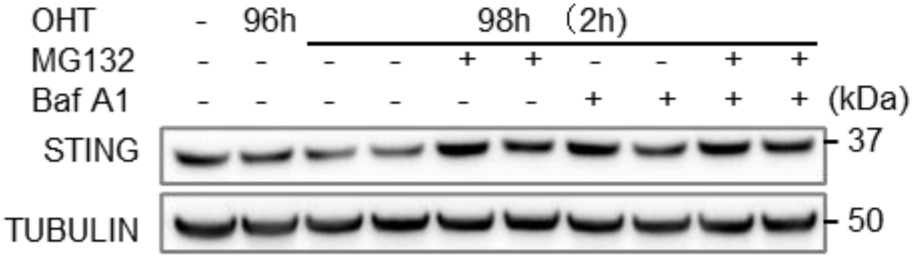
STING protein expression in MEFs following various treatments. Western blot analysis of STING protein expression in MEFs treated with OHT for 96 hours, followed by stimulation with MG132 and Baf A1 for an additional 2 hours.

**Figure S4.**
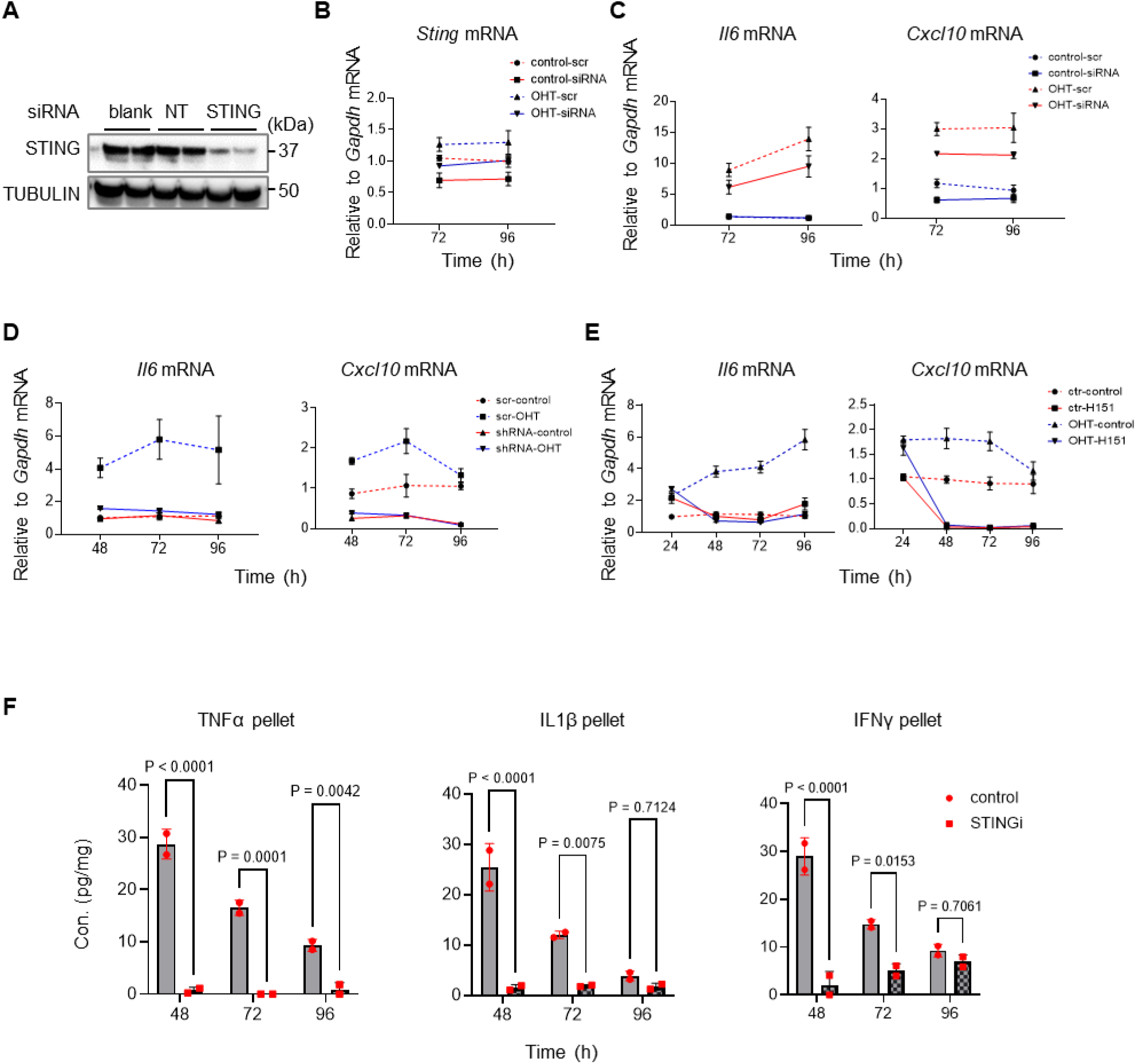
STING is essential for the activation of the inflammatory response in TRF1-deficient MEFs. (A) Western blot analysis confirming STING knockdown in MEFs transfected with either non-targeting control or STING siRNA for 72 hours. (B) MEFs were treated with OHT for 24 hours, followed by STING siRNA transfection for 48–72 hours. *Sting* mRNA levels were measured by RT-qPCR. (C–E) Expression levels of *Il6* and *Cxcl10* mRNA were analyzed at various time points following treatment with *Sting* siRNA (C), *Sting* shRNA (D), or STING inhibitor (E), in the presence or absence of OHT. (F) TNFα, IL1β, and IFNγ protein levels were measured by cytokine array in cell pellets from OHT-treated MEFs for indicated times and normalized to total protein concentration in the cell pellet. H-151 was administered 24 hours post-initiation of OHT treatment. P values were calculated using two-way ANOVA for multiple comparisons. Data are presented as mean ± SD from a representative experiment out of three independent experiments.

**Table S1:**
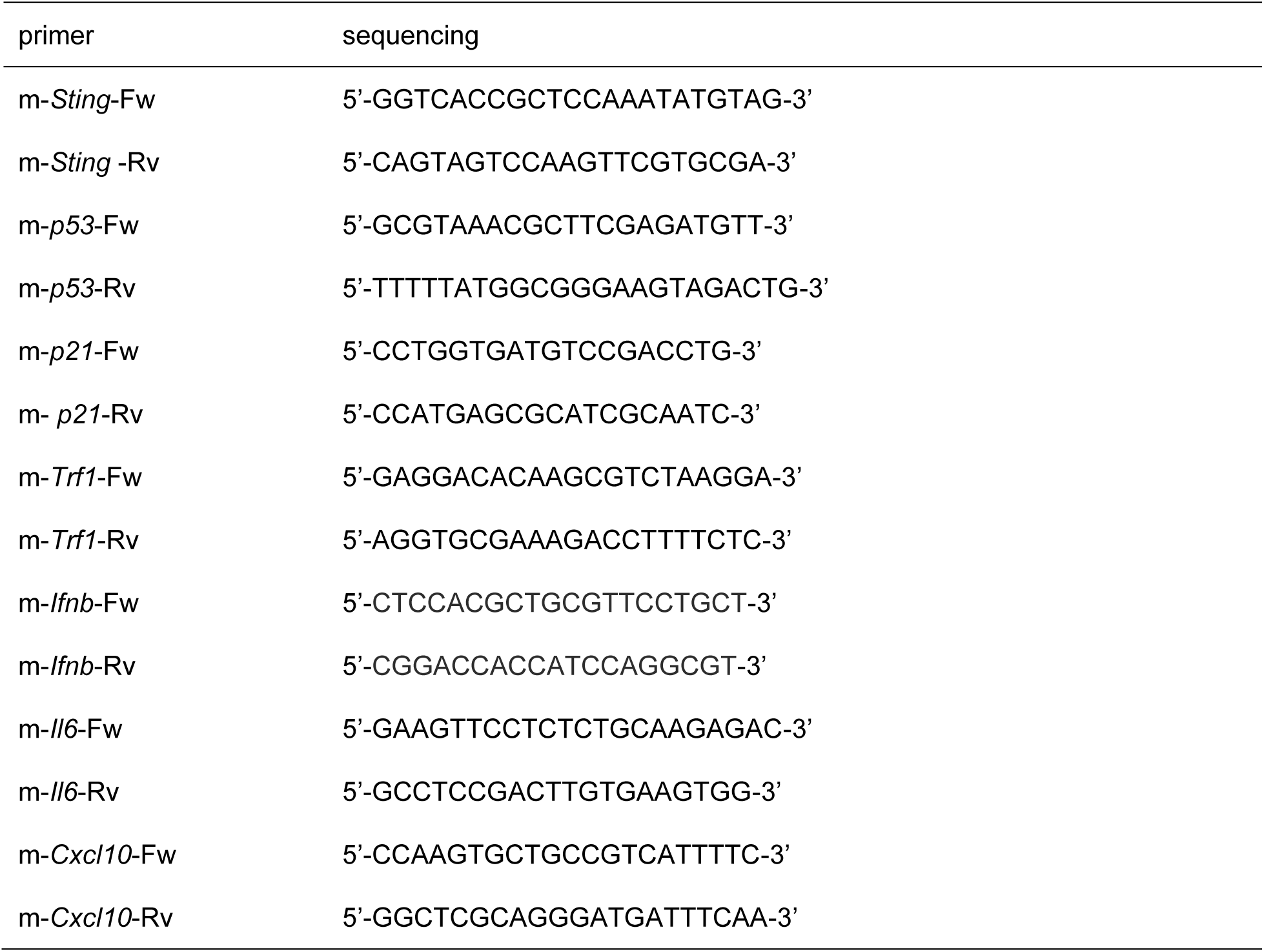
The primer sequences used for qPCR

